# Sequence conservation need not imply purifying selection: evidence from mammalian stop codon usage

**DOI:** 10.1101/2022.03.02.482615

**Authors:** Alexander T. Ho, Laurence D. Hurst

## Abstract

The assumption that conservation of sequence implies the action of purifying selection is central to diverse methodologies to infer functional importance. In mammals, however, GC-biased gene conversion (gBGC), a meiotic mismatch repair bias strongly favouring GC over AT, can in principle mimic the action of selection. As mutation is GC→AT biased, to demonstrate that gBGC does indeed cause false signals requires confidence that an AT-rich residue is selectively optimal compared to its more GC-rich allele, while showing also that the GC-rich alternative is conserved. We propose that mammalian stop codon evolution provides a robust test case. Although in most taxa TAA is the optimal stop codon, TGA is both abundant and conserved in mammalian genomes. We show that this mammalian exceptionalism is well explained by gBGC mimicking purifying selection and that TAA is the selectively optimal codon. Supportive of gBGC, we observe (i) TGA usage trends are consistent at the focal stop and elsewhere (in UTR sequences), (ii) that higher TGA usage and higher TAA→TGA substitution rates are predicted by high recombination rate and (iii) across species the difference in TAA <-> TGA rates between GC rich and GC poor genes is largest in genomes that possess higher between-gene GC variation. TAA optimality is supported both by enrichment in highly expressed genes and trends associated with effective population size. High TGA usage and high TAA→TGA rates in mammals are thus consistent with gBGC’s predicted ability to “drive” deleterious mutations and supports the hypothesis that sequence conservation need not be indicative of purifying selection. A general trend for GC-rich trinucleotides to reside at frequencies far above their mutational equilibrium in high recombining domains supports generality of these results.

## Introduction

If at a given site in DNA a mutation appears in a population and is eliminated by selection owing to its deleterious effects, the site in question will tend to be more conserved between species than comparable neutrally evolving sequence. This simple logic underpins the notion that the functionality of sequence can be inferred from its degree of conservation – for discussion see Ponting (1). It is explicit in, for example, molecular evolutionary tests for purifying selection (e.g. Ka/Ks test (2–5)), attempts to identify sites prone to disease-causing mutations (6, 7), and estimates of the proportion of DNA within a genome that is “functional” (8).

These methods assume, however, that no force other than selection can deterministically act to alter the frequency of extant alleles. Over the past two decades GC-biased gene conversion has been established as a potentially important influence on allele frequencies (9), mimicking selection (10–12). The process of gBGC results from a repair bias favouring G/C alleles over A/T alleles during GC:AT mismatch repair in a (commonly assumed to be meiotic) heteroduplex (13, 14). In humans, at non-crossover gene conversion events 67.6% of GC:AT mismatches favour the GC allele (15). It is probably as a consequence of this bias, coupled with the regionalisation of recombination domains over extended time periods, that mammals, alongside birds and possibly other amniotes (16), have genomes with large (> 300Mb) blocks of relatively homogeneous higher or lower GC content (isochores) (10, 11, 17). Importantly, assuming consistency of local recombination rates over evolutionary time and a correlation between crossover rates and non-crossover rates (18), gBGC also can explain the relatively strong correlation between GC content of these blocks and local recombination rates in mammals (19–22) (but see also 23, 24). That the correlation is stronger with male meiotic events than female ones is taken as evidence that the trends cannot be explained by selection with reduced Hill-Robertson interference in domains of high recombination (22). Consistent with such models, SNP analysis reveals the predicted fixation bias for AT→GC mutations in GC rich domains, even after allowing for non-equilibrium GC content (25, 26).

While the human conversion bias is strong, defining the expected impact of gBGC on the human genome is not trivial. For example, in any given generation, the net effect of bias is a function of the length of the relevant conversion tracts, the commonality of AT:GC mismatches within the tracts and the rate of initiation of such tracts. Williams et al. (18) estimate a rate in human non-crossover events (where there is the strong GC:AT bias) of 5.9 × 10^−6^ per bp per generation. More generally, Glemin et al. (27) estimate that the net effect on substitutions is on average in the nearly-neutral area. However, as recombination occurs primarily within recombination hotspots ∼2% of the human genome is subject to strong gBGC in any generation (27). Over the longer term as the location of recombination hotspots evolves rapidly, they predict that a large fraction of the genome is affected by short episodes of strong gBGC (27). Galtier (28) estimates that ∼60% of all synonymous AT→GC substitutions are influenced by gBGC.

Strong gene conversion is, however, not phylogenetically universal. In the best resolved instance, yeast, where meiotic tetrads can be directly studied, the bias is extremely weak at best. The highest estimates suggests that the GC-allele is the donor allele in 50.62% of cases (11, 29). Further analysis report a lesser bias (30), with a further large study reporting weak bias in the opposite direction (31). Meta-analysis of over 100,000 GC:AT mismatch resolutions in *Saccharomyces cerevisiae* determined a net segregation of 50.03%, only just in favour of the GC alleles and not significantly different from 50:50 segregation (31). To date strong conversion has been observed in only a few taxa (31), mammals (11) and birds (32, 33) being the two well-described exceptions, though weaker and non-regionalised gBGC is suspected in many taxa (21).

In terms of the population genetics influence, the action of gBGC is directly comparable to meiotic drive (alias segregation distortion) (34). In this sense gBGC may be said to “drive” alleles. In turn, such drive can mimic positive selection (35). Importantly, it has previously been noted that gBGC can (and in birds and mammals regularly does) create false signals of positive selection by promoting the spread from rare to common of AT→GC mutations (12, 36–40). However, as is implicit in all such models (41), gBGC could also mimic the action of purifying selection. A GC allele at fixation mutating to a selectively advantageous AT allele would be forced by gBGC to eliminate the AT allele, causing conservation of the deleterious GC allele.

Mimicry of positive selection owing to gBGC in mammals is thought to be common and, to date, analyses have focused on the substitional process, rather than the conservation process (12, 36–40). We are aware of no clear example of gBGC causing false signals of purifying selection. A core difficulty is finding a circumstance where gBGC makes predictions different from those of mutation bias and selectionist models. Differentiating between the effects of gBGC and mutation bias tends to be relatively straightforward as mutation is near-universally GC→AT biased (42–46), while gBGC is biased in the opposite direction. More problematic is the possibility that the GC state is also the selectively optimal state. If so, then both gBGC and selection make the same predictions of conservation of GC and covariation with the recombination rate. Given Bengtsson’s argument, that gBGC may be biased in this direction to counter a deleterious GC→AT biased mutational process (47), it may well be unusual to have the selectively optimal state being promoted by mutation bias but not by gBGC. Indeed, in Drosophila, for example, “optimal” codons tend to end G or C (48). Codon optimality may also not be adequate to define the direction of selection, however, as such selection may also be contingent on the overall GC-richness of the sequence (owing to RNA structure effects (41)). Thus, the core difficulty to establish gBGC as a cause of false signals of purifying selection and cause conservation of deleterious alleles is to identify a case where we can have confidence (and independently verify) that the AT state is selectively optimal compared to its GC-richer allele.

Here we suggest that mammalian stop codon usage may provide an exceptional test case. Across all domains of life the three stop codons, TAA, TGA and TAG, are not used equally (49), with TAA being commonly, if not universally, selectively favoured (49). This is probably owing in large part to selective avoidance of translational read-through (TR). During TR, the stop codon is missed by its cognate release factor (50) due to the mis-binding of a near-cognate tRNA (51, 52), leading to the erroneous translation of the 3’ UTR and the generation of potentially-deleterious protein products (53). Each stop codon has a distinct intrinsic error rate such that TGA>TAG>TAA in bacteria (54–59) and eukaryotes (55, 60) (including humans (61)). TR rate reduction in any given gene might thus be achieved by selection for TAA.

Evocation of such selection presumes that TR is usually deleterious (62, 63). This is likely as the formation of C-terminal extensions cause energetic wastage (64) as well as problems with protein stability (65–67), aggregation (68, 69), and localisation (70, 71). Alternatively, in the absence of another 3’ in-frame stop codon, both the read-through transcript and nascent protein are likely to be degraded when the translational machinery reaches the polyA+ tail (72, 73). In addition to reducing TR costs, TAA also has several other benefits: there may be selection for fast release of the ribosome to prevent ribosomal traffic jams (74) and it is robust to two mistranscription events (TAA→TGA, TAA→TAG) while the two other stop codons are resilient to just one (TGA→TAA, TAG→TAA).

It is then noteworthy that stop codon usage in mammals is different to that seen elsewhere (49, 75): TGA is more often conserved than TAA (76) and, unusually, the substitution rate of TAA→TGA is higher than the reverse (49). Despite the fact that in humans TAA is disproportionately employed in highly expressed genes (77), this signal of conservation has been interpreted as evidence that purifying selection is operating to preserve TGA in mammals (76). Gene conversion would however oppose fixation of TGA→TAA mutations (while also favouring TAA→TGA) and hence mimic purifying selection on TGA, even if selection were operating in the opposite direction. Biased gene conversion, thought to be especially influential in humans (15), could thus resolve the exceptionalism of TGA conservation in mammals.

Here we evaluate this suggestion. Duret and Galtier (11) provide a series of tests for differentiating gBGC from selection noting that the trend to the higher GC state should be correlated with recombination and common to all sites regardless of functional status. We consider several analyses to examine these predictions finding all to be robustly supported. However, to be confident that TAA underusage at the focal stop codon is indeed maladaptive, we also need confidence that TAA is the optimal stop codon. We consider several tests all of which support this. Finally, we resolve that complex mutational biases cannot fully explain the TAA/TGA usage trends and confirm a general pattern for GC-rich trinucleotides to reside at frequencies far above their mutational equilibria in GC-rich (high recombining) domains. The latter results are consistent with broadscale patterns of conservation of GC-rich residues owing to gBGC. The same analysis resolves the trinucleotide usage in domains not likely to be subject to gBGC is as expected from a model of complex mutation bias. Indeed, these models predict higher TGA usage than TAG usage in these domains. However, different trinucleotides of same nucleotide content (as with TGA and TAG), have repeatable differences in the extent to which they are subject to fixation bias in GC rich isochores. The cause of these previously unknown complex fixation biases is unresolved.

## Results

### Bias towards TGA usage is also evident in the 5’ and 3’ UTR

The gBGC hypothesis predicts that, because the AT→GC bias in the mismatch repair process is non-specific to terminating stop codons, stop codon usage at the focal stop need not be greatly different to usage of the same trinucleotides seen elsewhere in the genome. To address this, we analyse “stop” codon usage at the focal termination site and in human 5’ and 3’ UTR sequences irrespective of reading frame. This controls for effects of transcription coupled mutational bias. A model supposing that TGA is optimal in mammals predicts the patterns of stop codon usage as a function of GC content should not be seen in 5’ and 3’ UTR sequence.

We first establish how intronic GC, as a proxy for isochore GC, covaries with stop codon usage at the focal termination codon. Consistent with the observations of Seoighe et al. (76) and Belinky et al. (49), we find TGA to be the most common stop in the primate lineage (Fig 1). Not only is TGA the most common stop, it also significantly and positively covaries with intronic GC content in humans when both metrics are calculated in 10% percentile bins (n = ∼1000 genes) (Spearman’s rank; p < 2.2 x 10^-16^, rho = 0.99, n = 10). TAG usage is also correlated with intronic GC content (Spearman’s rank; p = 0.0014, rho = 0.89, n = 10). TAA frequency is negatively correlated with intronic GC content Spearman’s rank; p < 2.2 x 10-16, rho = -0.99, n = 10). As predicted by a gBGC model, we see the same trends in non-coding sequences. TAA frequency is negatively correlated with intronic GC content in both 5’ and 3’ UTR sequence (Spearman’s rank; both p < 2.2 x 10^-16^, both rho = -0.99, n = 10). TGA is positively correlated with intronic GC content in both 5’ and 3’ UTR sequence (Spearman’s rank; both p < 2.2 x 10^-16^, both rho = 1, n = 10). TAG is uncorrelated with intronic GC content in both 5’ (Spearman’s rank; p = 0.10, rho = 0.55, n = 10) and 3’ UTR sequence (Spearman’s rank; p = 0.61, rho = 0.19, n = 10). Analysis on a gene-by-gene basis (instead of using binned data) using linear regression models supports these conclusions and the same trends in stop codon usage can be seen in intronic sequence against GC3 (GC3 being used in this circumstance as intronic stop usage predicted by intronic GC would be non-independent; S1 Table). This is strong evidence that the trends in canonical stop usage are approximately the same as the trends in stop usage outside of the canonical termination context.

**Fig 1.**
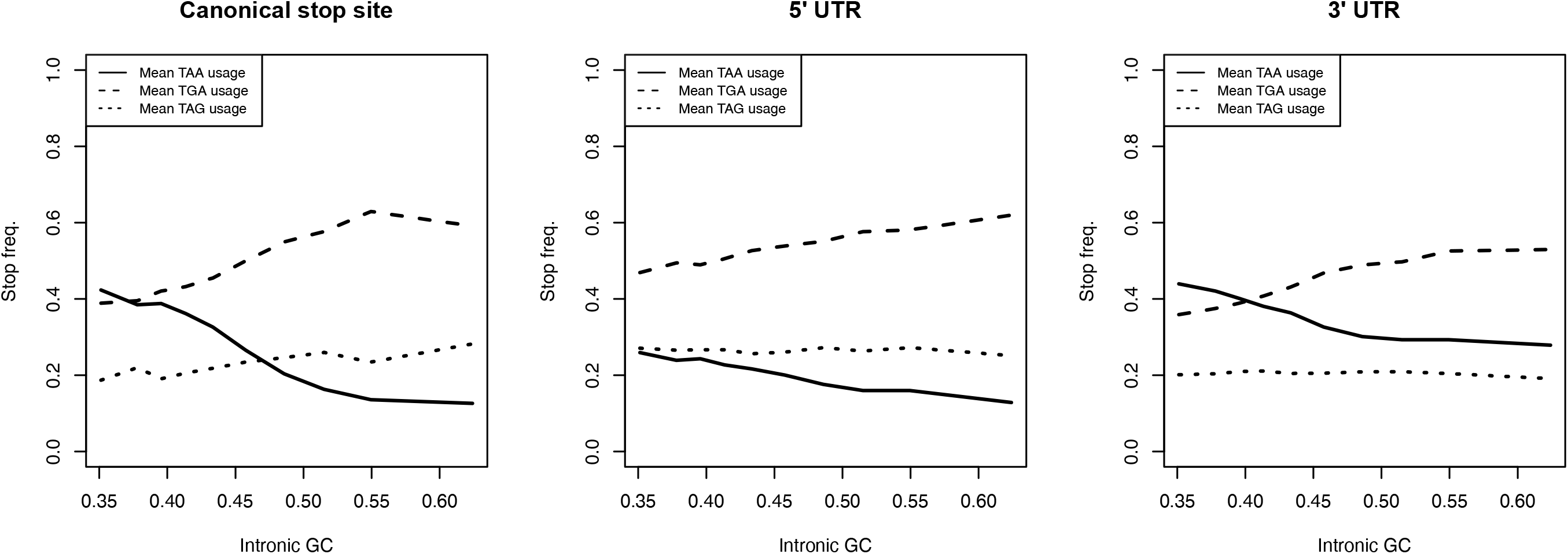
Stop codon frequencies (relative to the usage of all stops) at the canonical stop site, in the 5’ UTR, and in the 3’ UTR at ten equal sized bins of various intronic GC contents in the genome. TAA frequency is negatively correlated with intronic GC content in all three sequences (Spearman’s rank; all p < 2.2 x 10^-16^, all rho = -0.99, n = 10). TGA is positively correlated with intronic GC content in all three sequences (Spearman’s rank; all p < 2.2 x 10^-16^, rho = 0.99 for CDS, rho = 1 for both UTRs, n = 10). TAG usage is positively correlated with intronic GC content at the canonical stop site (Spearman’s rank; p = 0.0014, rho = 0.89, n = 10) but is uncorrelated with intronic GC content in both 5’ (Spearman’s rank; p = 0.10, rho = 0.55, n = 10) and 3’ UTR sequences (Spearman’s rank; p = 0.61, rho = 0.19, n = 10). Standard error bars calculated within each bin are extremely small and hence not shown.

### High TGA usage is strongly predicted by high recombination rate

Biased gene conversion can explain a strong correlation between the local recombination rate and substitution-derived GC* in primates (78, 79), GC* here being the predicted fixation bias determined equilibrium value rather than a non-equilibrium observed value. Similarly, such a model could predict high TGA usage in domains of high recombination. If TAA is optimal, the selection would not predict this as Hill-Robertson interference predicts more efficient selection with higher recombination rates.

To consider the effect of recombination on stop codon usage we consider both local instantaneous measures of recombination (from the HapMap 2 project, see methods) and broader scale analysis. The disadvantage of the former is that local recombination rates are not stationary over evolution time so current estimates need not reflect the past-history that influences stop codon usage. One problem with the latter is low samples size. Indeed, genome segments with consistently high recombination rates that could make for an ideal test are the pseudoautosomal regions (PAR1 and PAR2). However, there are few pseudoautosomal genes. As predicted by the gBGC model these regions have high GC content relative to the chromosome average, reportedly 48% in PAR1 compared to 39% in the rest of the X chromosome (80). In support of the gBGC model explaining high TGA usage, we also find that TGA is used much more often in PAR1 genes (71.4%, using one candidate transcript per gene annotated in this region) compared to the genome wide average (52.4%). Statistical comparison of TGA usage between these two values is, however, underpowered due to there being a low number of annotated genes which we may extract (n = 14).

A better “gross” scale analysis is to consider chromosome size as smaller chromosomes are associated with higher recombination rate per bp (21). As predicted by the gBGC model in the human genome, we find autosomal size (bp length) to be negatively associated with GC content (Spearman’s rank; p = 0.0078, rho = -0.56, n = 22) and TGA usage (Spearman’s rank; p = 0.0094, rho = -0.55, n = 22) (S1 Fig).

To test whether local recombination rate is predictive of stop codon usage in humans we employ logistic regression modelling considering all genes, using local recombination rate as the independent variable. Here we consider the recombination rate which for humans is valid as gBGC associated non-crossover and crossover events are highly correlated (18). We find that high recombination rate is significantly predictive of higher TGA usage (coefficient = 0.017, p = 0.023) and lower TAA usage (coefficient = -0.046, p = 1 x 10^-6^), these being the directions predicted by the gBGC hypothesis. Indeed, we find the same trends in non-CDS sequences when using linear models to predict trinucleotide frequencies as TAA, TGA, and TAG may appear more than once (unlike at the canonical stop). High recombination rate significantly predicts higher TGA trinucleotide frequency in the 5’ UTR (coefficient = 0.0032, p = 0.012), in the 3’ UTR (coefficient = 0.0053, p < 2.2 x 10^-16^), and in intronic sequence (coefficient = 0.0054, p < 2.2 x 10^-16^). It also significantly predicts lower TAA trinucleotide frequency in the 3’ UTR (coefficient = -0.0050, p = 3.5 x 10^-14^) and in intronic sequence (coefficient = -0.0043, p < 2.2 x 10^-16^), but not in the 5’ UTR where the regression coefficient is negative but not significant (coefficient = -0.0011, p = 0.28). These results are all consistent with gBGC promoting TGA over TAA in domains of high recombination both at the focal stop codon and elsewhere.

### Net flux to TGA stop codons is highest in GC rich and highly recombining genes

#### (i) Increased TAA→TGA substitution in GC-rich regions is common to mammalian and avian lineages, but not lineages that possess weak gBGC

The above considers observed patterns of usage. We can also consider evidence from recent substitution events. Here we consider flux, meaning the substitution rate from state A to state B (e.g. TAA→TGA) per occurrence of state A in the ancestral sequence. To calculate flux rates we consider species trios, assign an ancestral state to the internal node by maximum likelihood and calculate rates of change from this ancestral state to a derived state per incidence of the ancestral state. This is comparable to a prior method (49), excepting for our use of likelihood instead of parsimony.

The gBGC hypotheses predicts that TAA→TGA flux in the mammalian lineage should be highest in GC-rich isochores. More generally, it predicts that in species with gBGC strong and regionalised enough to cause high variation between genes in GC content, that the TAA→TGA flux should be especially accentuated in GC-rich domains. By contrast, species less influenced by gBGC should not show similar accentuation of TAA→TGA flux. We thus test whether the intragenomic difference in TAA<→TGA flux between the highest and lowest by GC is greater when the difference between the mean GC of the two partitions (high GC, low GC) is itself greater or when the intragenomic variance in GC is higher.

From the TAA→TGA and TGA→TAA flux rates, we may then adapt the formulae proposed by Long, Sung (44) to calculate TGA content from these flux rates alone, pTGA (see methods). This provides a single metric of the relative substitution rate between the two stop codons. This we do for the top (GC-rich) and bottom (GC-poor) 50% of genes by GC content, assayed by calculating the intronic GC content of each orthologue from one candidate species from the trio, to determine whether the TAA→TGA rate increases with GC pressure.

We calculate the difference in pTGA between GC-rich and GC-poor genes for 4 mammalian set of species trios (within primates, mice, dogs, and cows) and 4 non-mammalian species trios (birds, nematodes, flies, and plants) (see https://github.com/ath32/gBGC for species lists). To assay the extent of pTGA deviation we calculate (O-E)/E where O is pTGA of the GC rich set and E is that for the GC poor set of genes. To assign significance, we compare observed pTGA deviation scores to null simulations that calculate pTGA for two null groups of genes according to the net genomic TAA→TGA and TGA→TAA rates (see methods).

Consistent with the hypothesis that gBGC drives high TGA usage in GC rich isochores, pTGA is higher in GC-rich genes than GC-poor genes across the four mammalian lineages. The difference between the gene groups is greater than expected by chance in all four cases (primates: p = 0.014, dog: p = 0.040, cow: p < 0.0001, mouse: p < 0.0001). Of the non-mammalian lineages, pTGA in GC rich genes exceeds pTGA in GC poor genes in birds (p = 0.174), flies (p = 0.427), and nematodes (p = 0.231) but none of the observed differences are significantly different to null. Probably due to the selfing biology of Arabidopsis (81), pTGA is lower in GC-rich genes than GC-poor genes (nevertheless, p = 1 using the same test as the other lineages, Fig 2h).

**Fig 2.**
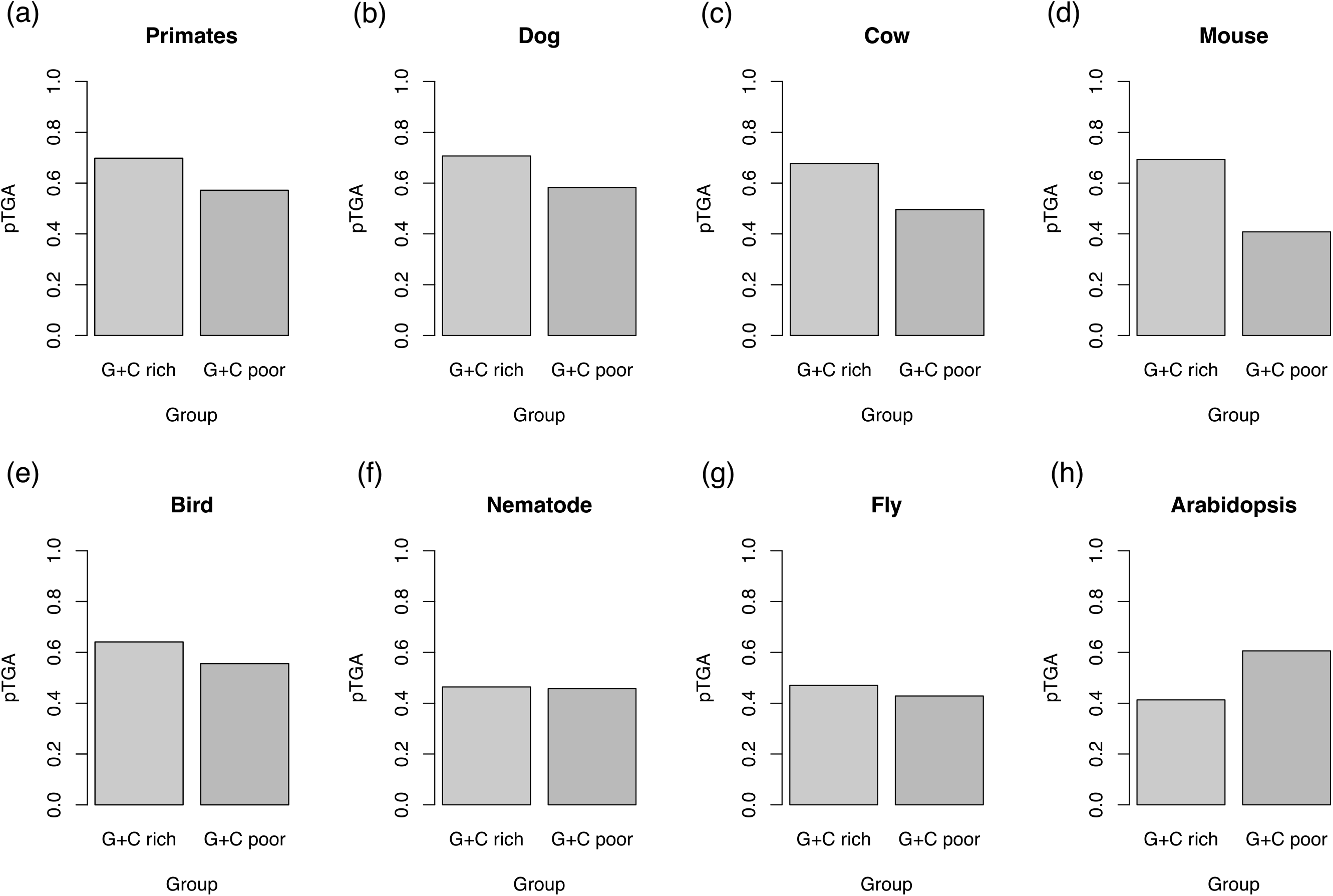
Predicted TGA usage (pTGA) derived from TAA→TGA and TGA→TAA flux for the top 50% of genes by GC content and bottom 50% of genes by GC content in four mammalian (a-d) and four non-mammalian (e-h) lineages. pTGA is calculated as 1/(1+(TGA→TAA/TAA→TGA)) and hence represents the balance between the two dominant stop codon flux events. Bootstrapped 95% confidence intervals are miniscule and hence not shown.

The prediction of the gBGC model is that the between-species variation in intragenomic flux difference should be predicted by the extent of GC variation within the genome. For this analysis, we calculate GC variation as the difference in mean intronic GC content between the two sets of genes analysed in Fig 2 and call this ΔGC. We also estimate the variance in GC3 between all genes. Consistent with the gBGC hypothesis for explaining TGA usage trends, analysing our eight lineages we find pTGA deviation is significantly correlated with both ΔGC (Spearman’s rank; p = 0.046, rho = 0.74, n = 8) and genomic variance in coding sequence GC3 (Spearman’s rank; p = 0.028, rho = 0.79, n = 8) (Fig 3).

**Fig 3.**
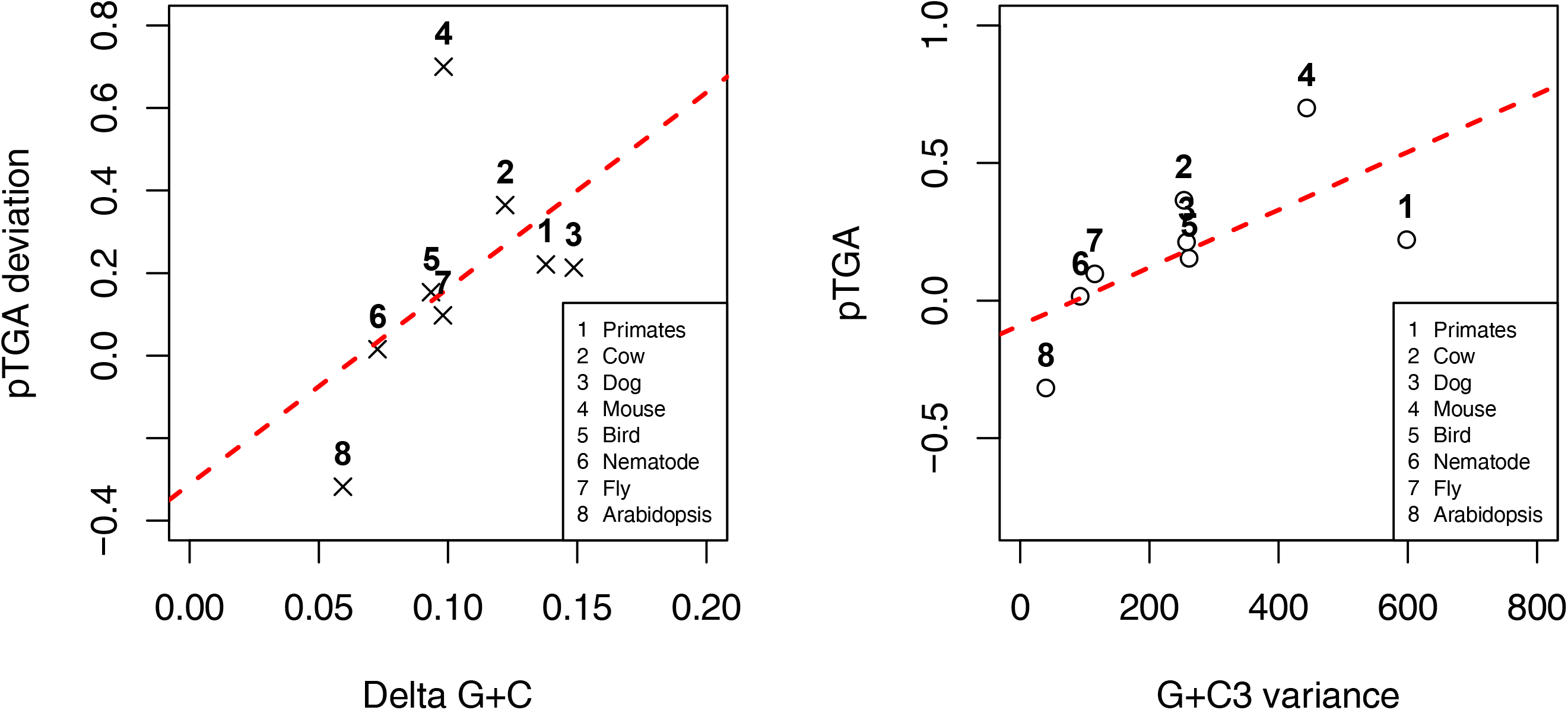
Predicted TGA usage (pTGA) deviation between the top 50% and bottom 50% of genes by GC content as a function of (a) the difference in GC content between the two gene bins, “delta GC”, and (b) coding sequence GC3 content variance across a sample of four mammalian and four non-mammalian lineages. pTGA is calculated as 1/(1+(TGA→TAA/TAA→TGA)) and hence represents the balance between the two dominant stop codon flux events. pTGA deviation is calculated as (O-E)/E where O is the pTGA score of GC rich genes and E is the pTGA score of GC poor genes. pTGA deviation is positively correlated with both delta GC (Spearman’s rank; p = 0.046, rho = 0.74, n = 8) and GC3 variance (Spearman’s rank; p = 0.028, rho = 0.79, n = 8). Bootstrapped error bars on the x and y axes are miniscule and not shown.

This suggests that species with pronounced TAA→TGA flux in their GC-rich domains (mammals) also tend to have more variation between their GC richest and GC poorest genes. Broadly these results accord with what is known about gBGC across these species. The (O-E)/E values are higher in mammals (primates = 0.221, cows = 0.365, dogs = 0.213, mice = 0.700) and birds (birds = 0154) than in invertebrates (nematodes = 0.015, fly = 0.097) and plants (Arabidopsis = -0.318). Birds are expected to resemble mammals as they too have pronounced gBGC (11, 82). However, small chromosomes and associated high recombination rates probably mean that most genes in birds are subject to considerable gBGC, it being notable that the predicted pTGA is high for both gene groups (Fig 2e). Non-isochore-containing genomes of invertebrates may possess AT→GC biased gene conversion, albeit with much weaker (31, 41, 83) or less regionalised effects. Arabidopsis being an almost obligate inbreeder is expected to be most affected by mutation bias and least affected by gBGC (81).

#### (ii) TAA→TGA flux is higher in highly recombining genes than lowly recombining genes

Just as gBGC predicts TAA→TGA flux to positively covary with GC content, as gBGC is coupled tightly to recombination it also predicts a positive relationship with recombination rate. To assess this, using data from the HapMap2 project, we first define highly recombining genes (HRGs) as the top 50% of genes by recombination rate and lowly recombining genes (LRGs) as the bottom 50%. Adapting our stop codon flux methodology, we then calculated the flux rates for TAA→TGA and TGA→TAA for HRGs and LRGs and used these rates to calculate pTGA for both groups (Fig 4). Significance was once again determined by comparing the observed pTGA deviation to those observed in null simulations that assume uniform genomic TAA→TGA and TGA→TAA rates. Consistent with the hypothesis that gBGC drives high TGA usage in highly recombining regions, pTGA is higher in HRGs than LRGs, (p=0.049). The pTGA deviation score between HRGs and LRGs is 0.172, slightly less than observed between GC rich and GC poor genes in the same genome (0.221).

**Fig 4.**
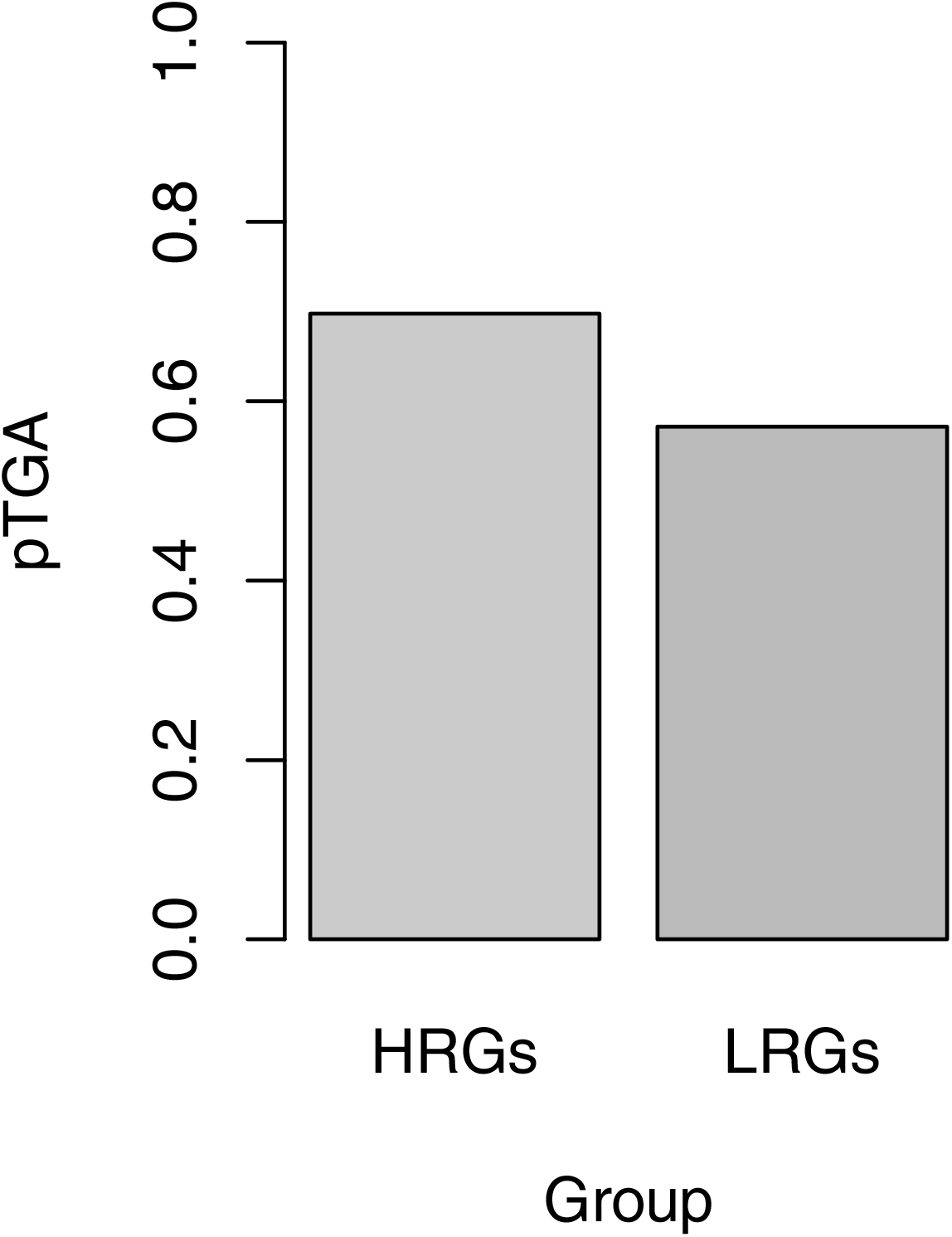
Predicted TGA usage (pTGA) derived from TAA→TGA and TGA→TAA flux for the top 50% of genes by recombination rate (HRGs) and bottom 50% of genes by recombination rate (LRGs) in the human genome. pTGA is calculated as 1/(1+(TGA→TAA/TAA→TGA)) and hence represents the balance between the two dominant stop codon flux events. Bootstrapped confidence intervals are miniscule and hence now shown.

### No evidence to support TGA optimality in eukaryotes

The evidence from non-termination sites supports the hypothesis that, whatever causes unusual TGA usage trends in most mammals, it cannot be explained by selection on the focal termination codon alone. Also, as predicted by the gBGC model, the TAA→TGA flux is stronger in domains of high GC/high recombination. Nonetheless, to have a case that gBGC acts against the direction of selection we need also to be able to confident that selection does not prefer TGA. Outside of the focal termination codon this is hard to assay but at the focal stop codon we can gather further evidence.

First, selection on any genic feature is classically assumed to predict that usage of that feature will be most common in highly expressed genes (62, 84, 85) as selection is strongest in highly expressed genes. Over-usage of “optimal” codons in highly expressed genes is a case in point (86, 87). In the current context, the opportunity for deleterious read-through (or other stop codon error) should scale linearly with the amount of protein product, so protein levels are a good metric for assaying strength of selection on the stop codon. Hence, if TGA usage were to be explained by selection, TGA usage is predicted to positively correlate with expression level. Prior data appeared to contradict this, suggesting that human highly expressed genes (HEGs, the opposite being LEGs) preferentially use TAA stop codons (77). However, possible covariation between expression level and GC content (88–90) could disturb the ability to make correct inference. We ask whether TAA or TGA are over employed in highly expressed genes controlling for GC content.

Second, the efficiency of both selection (91, 92) and gBGC (28, 32) are expected to vary with the effective population size (N_e_), both being more effective when N_e_ is high. The gBGC effect is however complicated by the fact that selection may also modify the effect of gBGC, reducing its impact if deleterious (28), such selection in turn also being dependent on N_e_. Most evidence suggests that gBGC is more influential when N_e_ is high (but see also (93)). However, we know the direction of gBGC, and it must act against TAA. Thus, across eukaryotes our expectation is that if TAA is optimal (and gBGC relatively less important), its usage will increase with N_e_. However, if gBGC is unexpectedly important outside of mammals or if TGA is optimal then TGA will increase with N_e_. We previously observed this not to be the case, with TAA increasing with N_e_ (94). However, the possibility remains that for lowly expressed genes TGA might be optimal and causing the focal termination codon trends (despite similar behaviour in 3’ UTR). We test this extension.

#### (i) High expression level strongly predicts high TAA usage controlling for GC

To test the predictive power of expression level on stop codon usage, we consider a series of logistic regression models. Each gene was assigned a 1 (present) or 0 (absent) in three different columns, TAA, TGA and TAG, depending on its stop codon identity. These scores were included as the dependent variable in several logistic regression models, with protein abundance (as a proxy for gene expression, for which we employ the natural log to promote a normal distribution) an independent predictor. We control for GC content by fitting multivariate models that include GC3 content (Table 1). Collinearity between GC content and protein abundance need not be a concern as the computed variance inflation factors are very low (less than 1.1 for all models).

**Table 1.** Results from multivariate logistic regression analysis that assess the extent to which gene expression and gene coding sequence GC content can predict stop codon usage in mammalian genes.

| Stop | Parameter | Primates |  |  | Dog |  |  | Cow |  |  | Mouse |  |  |
| --- | --- | --- | --- | --- | --- | --- | --- | --- | --- | --- | --- | --- | --- |
|  |  | Coef. | Std. Error | p-value | Coef. | Std. Error | p-value | Coef. | Std. Error | p-value | Coef. | Std. Error | p-value |
| TAA | Log(PxAbundance) | 0.023 | 0.007 | 6E-4 | 0.071 | 0.014 | 4E-7 | 0.071 | 0.008 | 2E-16 | 0.013 | 0.006 | 0.039 |
|  | GC3 | -0.038 | 0.001 | 2E-16 | -0.033 | 0.002 | 2E-16 | -0.040 | 0.002 | 2E-16 | -0.034 | 0.002 | 2E-16 |
| TGA | Log(PxAbundance) | -0.015 | 0.006 | 0.009 | -0.039 | 0.012 | 0.001 | -0.050 | 0.007 | 3E-13 | -0.006 | 0.005 | 0.273 |
|  | GC3 | 0.019 | 0.001 | 2E-16 | 0.018 | 0.002 | 2E-16 | 0.022 | 0.001 | 2E-16 | 0.019 | 0.001 | 2E-16 |

Consistent with prior observations of stop codon covariance with GC content (77, 95), we find TAA usage to be negatively (indicated by the sign of the coefficient), and TGA to be positively, correlated with GC3 in all four species trios tested. By the same coefficient analysis, we find that high TAA stop codon usage is predicted by high expression level in all four mammalian lineages (77), contra to the possibility that TGA has become the favoured stop codon in mammals. Both protein abundance and GC3 are consistently significant predictors of stop codon usage in our three mammalian lineages. In 8/8 models, the coefficients of protein abundance are consistent with TAA preference over TGA in highly expressed genes. Assuming that gene expression levels in orthologous genes are stable, stop codon usage reliably informs us of the stop that is preferred by selection.

#### (ii) Across taxa, lowly expressed genes also prefer TAA over TGA

While the above analyses provides support for the hypothesis that TAA, and not TGA, is preferred in highly expressed genes there is however, a further possibility, namely, that while TAA may well be preferred by HEGs, TGA may be optimal in LEGs. If this were to be the case, TGA might increase in genome-wide usage if most genes are not “highly” expressed. This we test by phylogenetically generalized least squares (PGLS) regression analysis that compares TGA enrichment (at the primary stop codon compared to downstream, to remove any GC covariance) in LEGs to effective population size (N_e_) for several eukaryotic species controlling for phylogenetic topology (see PGLS in methods).

We find N_e_ to be a significant negative predictor of TGA enrichment in LEGs (PGLS; estimate = -0.060, p = 0.012). By contrast, TAA enrichment in LEGs is positively, if not significantly, associated with N_e_ (PGLS; estimate = 0.073, p = 0.078). When we consider HEGs, N_e_ positively and significantly correlates with TAA enrichment (PGLS; estimate = 0.059, p = 0.0014) but is negatively, if not significantly, associated with TGA enrichment (PGLS; estimate = -0.044, p = 0.17). These results are not consistent with a selective preference for TGA stop codons at any expression level. These same results also indicate that gBGC is not an important force in most of the species examined as gBGC should also be more influential when N_e_ is high and force increased usage of TGA (32).

### TAA→TGA flux cannot be explained by mutation bias in humans

The above evidence indicates that whatever causes TGA conservation it is neither specific to the termination site nor explained by selection for termination efficiency at the termination site. In principle the trends we have seen could be explained by mutation bias. However, mutation bias tends to be GC→AT biased so should favour TAA not TGA (42–46). Nonetheless the possibility remains either that some more complex *k*-mer bias might exist or that mutation bias varies by isochore. Indeed, nucleotide pools can vary through the cell cycle potentially altering local mutation bias (96). Moreover, CpG to TpG rates are high in humans (97–100) and thus creation of new stop codons away from the focal stop (e.g. within 3’ UTR) via CpGA to TpGA could be common. We could imagine for example that focal stop codons commonly mutate to a sense codon this being rescued by a 3’ UTR pre-existing stop. If so, stop codon usage could be determined by mutational processes away from the focal termination codon. The same model does not however predict TAA→TGA flux at orthologous termination sites. That CpG deamination rate may also correlate negatively with GC content (100) also renders this an unlikely explanation.

We consider the relative rates of human germline *de novo* mutations derived from family trio data (101). From the mutation rate of each class of mutational event we calculate rates per occurrence of the ancestral nucleotide and generate a mutational matrix. From this we calculate the neutral equilibrium frequencies of all nucleotides (denoted N*), dinucleotides, or codons (see methods). From N* predictions we may predict the equilibrium GC frequency (GC*). Under the assumption that nucleotide contents are stationary, deviation of the observed nucleotide content from predicted equilibrium provides an indication of the direction of any fixation bias (44). However, equilibrium status is disputed (102) and the predicted equilibrium can vary with complexity of the mutational model (mono-nucleotide, di-nucleotide etc).

Consistent with previous analyses (42–46), from a dataset of 108,778 observed *de novo* mutations we find an overall GC→AT skewed mutational profile that hence fails to predict observed stop usage (S2 Table). Might, however, variation in mutation bias between isochores explain increasing usage of TGA and decreasing usage of TAA as domains become more GC-rich? To assay whether the above mutational profile covaries with intronic GC in a similar way to stop codon flux, we first repeat the above analysis for mutations found in different isochore GC contents (see also Smith et al. Smith, Arndt (46)). For each mononucleotide change, the local GC content (10kb window) was calculated. Mutations were then ordered by GC and split into 10% percentile bins of equal size (∼10,000 mutations each). From each of these bins and their associated mutational spectra and nucleotide contents, we recalculate GC* and TGA* (Fig 5, orange points). We find our GC* and TGA* predictions for each bin to be consistent between isochores of different GC content, indicating that mutation bias is not driving the trends we see in TGA usage nor TAA→TGA stop codon flux – and see also Smith et al. Smith, Arndt (46). If anything, mutation bias is increasingly GC→AT biased at high GC as the local GC content around de novo mutations is negatively correlated with their predicted GC* (Spearman’s rank; p = 0.024, rho = -0.72, n = 10) and TGA* (Spearman’s rank; p = 0.035, rho = -0.68, n = 10).

**Fig 5.**
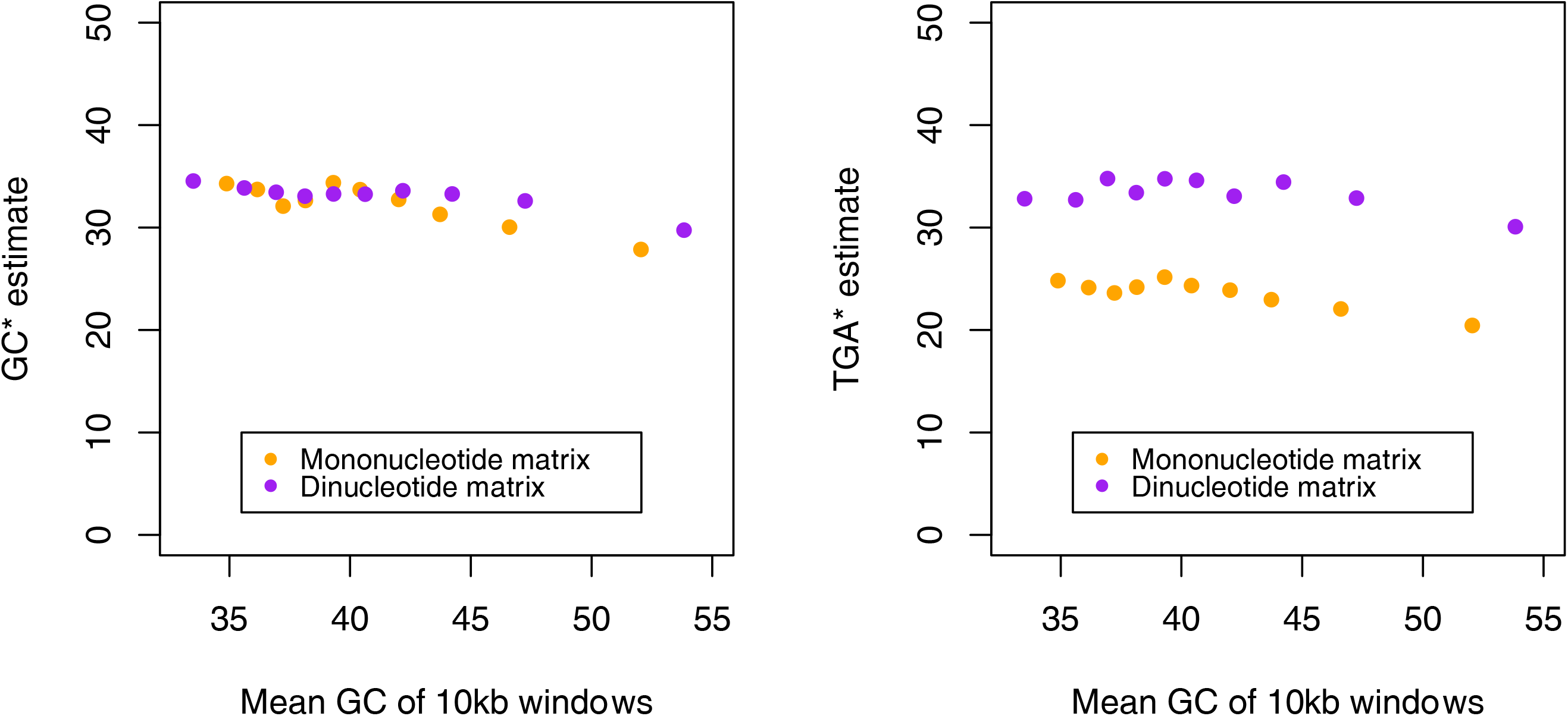
Predicted GC equilibrium (GC*) and relative TGA equilibrium (TGA*) frequencies across isochore GC contents derived from mononucleotide (orange) and dinucleotide (purple) mutational matrices. Standard deviations for the datapoints are minuscule and hence error bars are not shown (∼0.5% for mononucleotide estimates of TGA* and GC*, ∼0.5% for dinucleotide estimates of TGA* and ∼0.1 for dinucleotide estimates of GC*).

The above approach makes no allowances for more complex dinucleotide effects nor the possibility that some stop codons might be generated by mutations within CDS or within 3’ UTR sequences when the focal stop mutates. Given that there is hypermutability at CpG residues, leading to TpG residues (97–100) that are likely to affect the mutation-drift equilibrium frequency of TGA, we expand our analysis to consider the 16×16 dinucleotide mutational matrix. We also apply a model in which we generate null sequences from the equilibrium mutational matrix in a Markov process, hence allowing for within UTR mutational events. We consider the relative frequencies of the three stop codons in such sequence and how they vary by local GC. Consistent with the mononucleotide results, we find dinucleotide-derived GC* and TGA* to be lower than observed in the genome (40.9% and 52.4% respectively) and, importantly, flat across GC contents (Fig 5, purple points). While TGA* derived from the dinucleotide matrix exceeds TGA* derived from the mononucleotide matrix this is probably as a consequence of permitting CpG hypermutation generating potentially premature stop codons. We conclude that the absence of evidence for increasing GC* with GC content strongly argues against mutation bias as an explanation for higher TAA→TGA flux and higher TGA usage in GC-rich isochores.

### Mutation bias predicts trinucleotide usage in GC poor domains and TAG rarity

Above we have generated a mutational expectation for all trinucleotides but focused on TGA. This allows us to ask a series of further questions. For example, for all trinucleotides might a mutational null match what we see in GC poor domains, as expected if these are less subject to gBGC? In addition, can mutation explain any trends in stop codon usage in GC poor domains, for example the observation that TAG is underused compared with TGA?

We find that observed trinucleotide frequencies from GC poor sequences (the bottom 20% of genes by GC content) are accurately predicted by a GC poor mutational matrix (derived from the bottom 20% of de novo mutations by surrounding 10kb GC content) for all sequence that isn’t CDS (r^2^ > 0.9; Figure 6). This strongly supports the hypothesis that mutation bias alone may explain trinucleotide trends in GC poor domains outside of the coding context. In addition, while one can always consider more complex *k*-mer dependent mutational models, our extension from dinucleotides rates appears to be robust. Importantly, in such GC poor isochores TAG equilibrium is lower than TGA equilibrium (S2 Fig). This indicates mutation bias operates differently on the two, going some way to explain why TAG and TGA behave differently.

**Figure 6.**
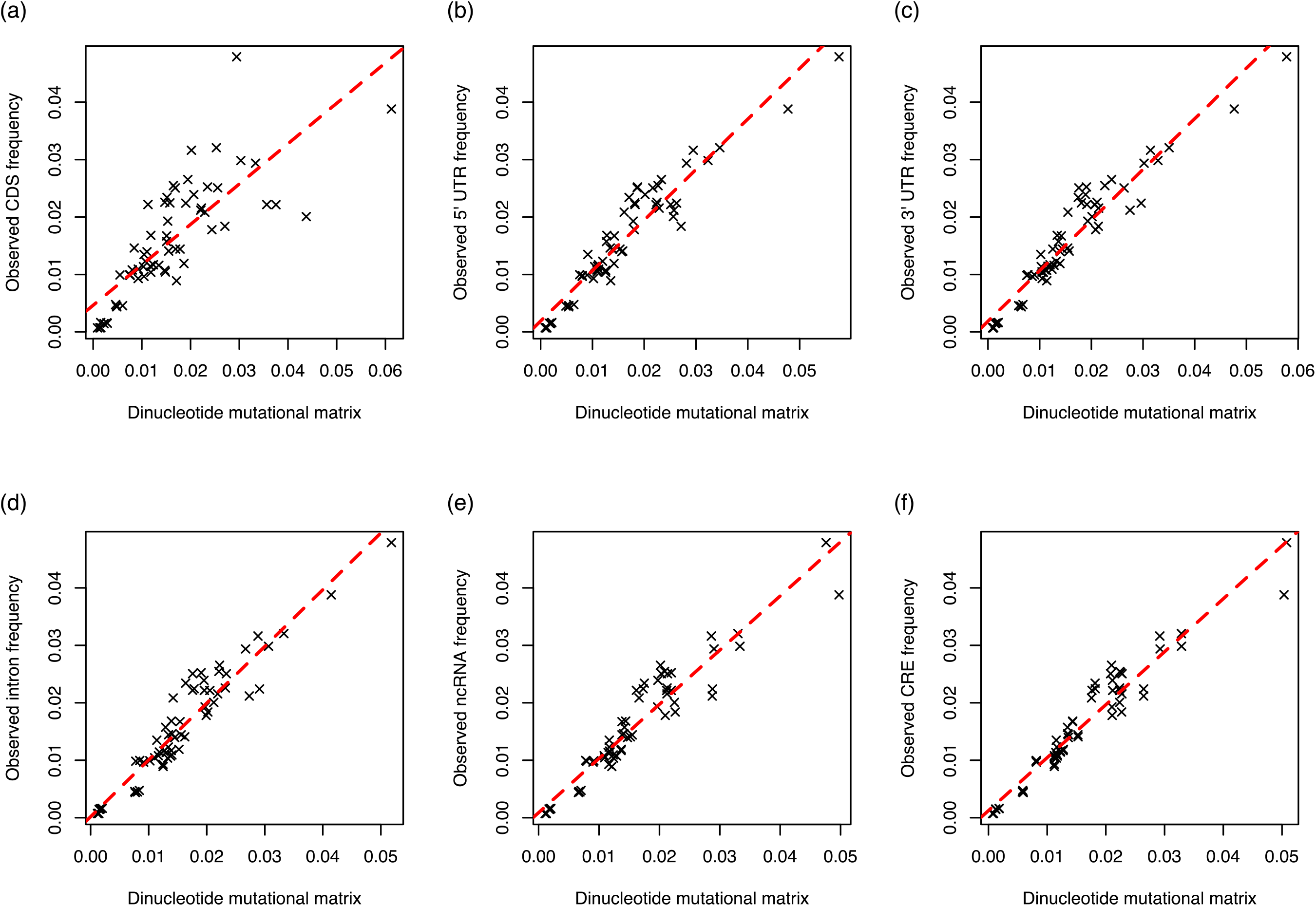
Observed (a) CDS, (b) 5’ UTR, (c) 3’ UTR, (d) intronic, (e) ncRNA, (f) cis regulatory element (CRE) trinucleotide frequencies as a function of the expected frequencies of the same trinucleotides derived from a dinucleotide mutational matrix. Expected frequencies were calculated simulated DNA sequences derived from dinucleotide equilibrium frequencies. Dinucleotide frequencies were calculated from a sample of de novo mutations taking place in the bottom 20% of sequences by GC content to avoid potential GC-coupled fixation biases. Expected frequencies accurately predict what is seen in real CDS sequence (linear regression; p = 7.7 x 10^-15^, adjusted r^2^ = 0.62), 5’ UTR sequence (linear regression; p < 2.2 x 10^-16^, adjusted r^2^ = 0.90), 3’ UTR sequence (linear regression; p < 2.2 x 10^-16^, adjusted r^2^ = 0.91), intronic sequence (linear regression; p < 2.2 x 10^-16^, adjusted r^2^ = 0.90), ncRNA sequence (linear regression; p < 2.2 x 10^-16^, adjusted r^2^ = 0.90), and CRE sequence (linear regression; p < 2.2 x 10^-16^, adjusted r^2^ = 0.93).

### gBGC predicts deviations from mutational expectations for all trinucleotides

The previous analysis suggests that in low GC domains *k*-mer trends are well predicted by mutation bias alone (Figure 6). By contrast, in GC rich domains, there exists a substitutional bias to TGA that is incompatible with mutation bias alone (Figure 5). Is the TAA→TGA fixation bias in high GC domains illustrative of a broader pattern? Were gBGC mimicking purifying selection we expect that GC rich trinucleotides should be most deviant from their mutational null in GC-rich domains. We hence extend the above analysis to consider the extent to which all trinucleotides deviate from mutational equilibrium as a function of their isochore of residence. In this instance, however, we cannot be confident that the GC-rich residue is selectively deleterious (as with TGA). Moreover, even when optimal codons are known to be GC ending selection at exon ends can commonly be in the opposite direction to enable accurate splicing (103), adding complexity.

Using mutational profiles from the relevant isochore, we calculate trinucleotide frequencies that represent our mutational null and compare these to observed trinucleotide frequencies in the genome. To test the hypothesis that a fixation “boost” in GC-rich isochores acts differently on GC-rich trinucleotides, we calculate a fixation boost metric. Specifically, we first calculate a (Observed-Expected)/Expected score for the top 20% of sequences by GC content, where expected is the mutational equilibrium frequency derived from the top 20% of de novo mutations assaying their surrounding 10kb GC content. This metric we term deviation 1, or *D1* for short. We then repeat this for the bottom 20% of sequences by GC content using their equivalent set of de novo mutations, receiving *D2*. Given the above results (Fig 6), we expect the bottom 20% to be closest to mutation equilibrium, hence having a low *D2* score. By contrast if there is a GC-correlated fixation bias, *D1* should be high for the GC-rich trinucleotides. We thus consider, for each trinucleotide, the difference between *D1* and *D2* values, this reflecting the shift in fixation process associated with domains of high GC. Using this metric, trinucleotides may be ranked by the “boost” they receive from GC-coupled fixation bias within their GC class. Thus, we classify all trinucleotides into one of four classes by GC% (0%, 33%, 66%, 100%). Within the zero class are trinucleotides with no G or C (e.g., AAA, ATA, TTA) and within the 100% class by contrast are those with no A or T (e.g., GGC, GGG), for example.

We find that the more GC-rich the class of trinucleotides the more they exceed their mutational equilibrium in high GC isochores (0% < 33% < 66% < 100%) (for statistics see Figure 7). This strongly supports the notion that the trinucleotide content of isochores derives from a fixation bias, rather than mutation bias, favouring GC residues, as gBGC would predict. More generally then, we have strong reason to suspect the gBGC-mediated fixation bias causes false signals of purifying selection at GC-rich residues in GC-rich isochores that extend far beyond the specific context of TAA→TGA flux.

**Figure 7.**
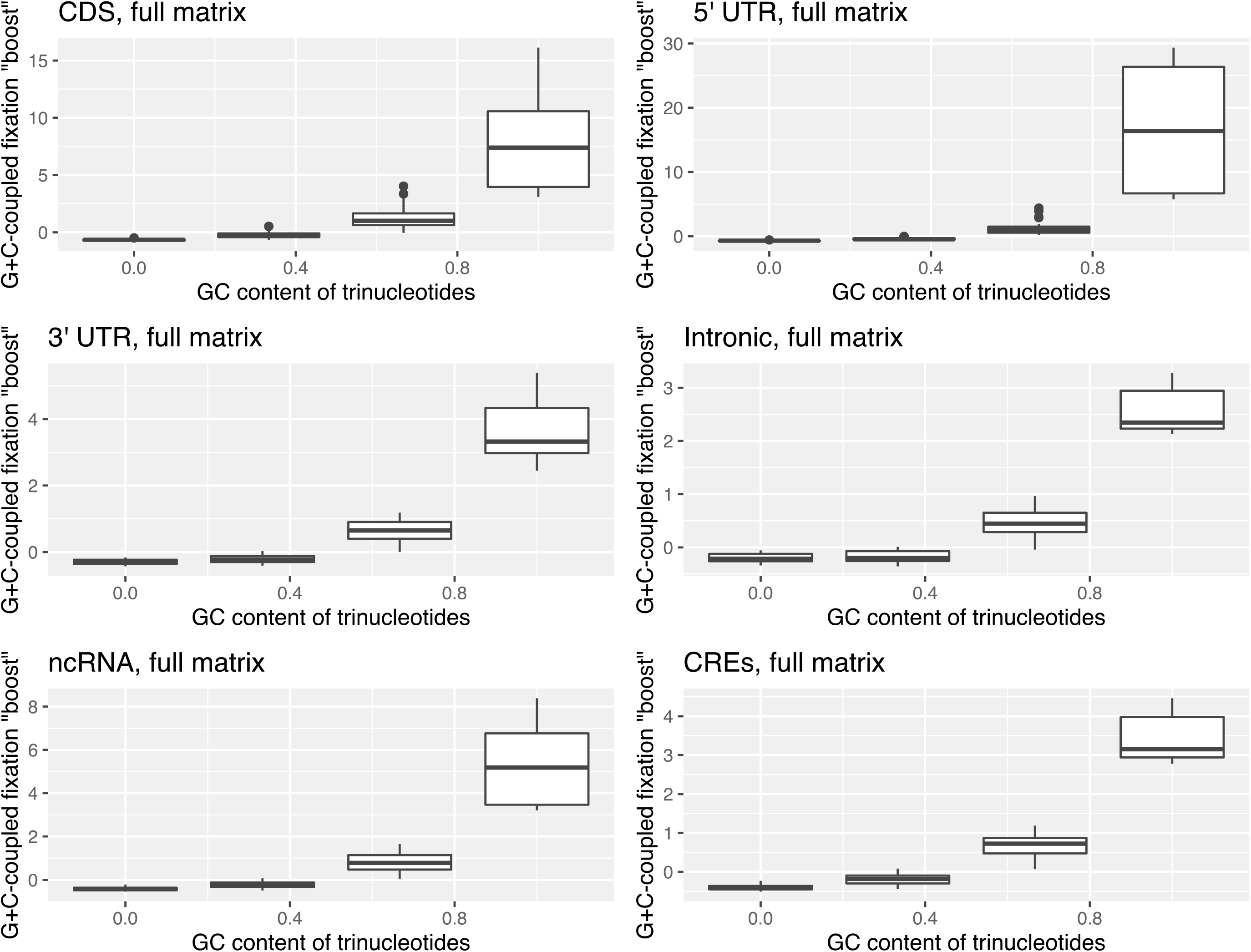
Deviation scores, (O-E)/E, describing the difference in GC-coupled fixation “boost” for the four GC classes of trinucleotides. Deviation between fixed and mutational equilibrium frequencies for each trinucleotide in the top 20% of sequences by GC content, D1, was calculated as (Observed-Expected)/Expected, where expected is the mutational equilibrium frequency. This was repeated for the bottom 20% of sequences by GC content to receive D2. As we predict GC-rich sequences to be subjected to stronger biased gene conversion, we predict D1 > D2. To compare D1 and D2 we once again calculate (Observed- Expected)/Expected, which we dub the GC-coupled fixation “boost”. In all sequences, GC content is positively correlated with this “boost” metric (Spearman’s rank; all p < 2.2 x 10 ^-16^; rho = 0.92 in CDS, rho = 0.94 in 5’ UTR, rho = 0.90 in 3’ UTR, rho = 0.87 in introns, rho = 0.92 in ncRNA, rho = 0.93 in CREs, n = 64 in all tests).

We assess this possibility a second way by considering flux between all two-fold synonymous codon pairs, all ending G:A or C:T, in genes of increasing recombination rate. Considering all two-fold synonymous codon pairs en masse, we find that the flux to the GC-rich codons are most strongly favoured at high recombination rates, consistent with possible gBGC action (S3 Fig). Before Bonferroni correction, this is true for 10 of the 12 two-fold synonymous codon pairs individually (Binomial test with null probability = 0.5; p = 0.039). This too is supportive of a gBGC-mediated fixation bias that is much more general than the stop codon example. Unlike with TAA and TGA flux, however, we can’t in these examples be sure which (if either) is the selectively optimal state. The two exceptions are Leucine (TTA<->TTG) and Glutamine (CAA<->CAG) where the ratio of flux increasing GC and decreasing GC is invariant to recombination rate (S4 Fig). That both CAA<->CAG and TAA<->TAG are unrepresentative of the more general trend is noteworthy.

### Trinucleotides have stereotypical fixation biases

We have observed that high TGA usage and high TAA→TGA fixation bias is especially common in GC-rich isochores, but TAG usage does not behave in the same way. Is this difference between two GC-matched trinucleotides particular to TAG and TGA? The CAA→CAG result would suggest not. We can address this by considering within GC-class variation in the fixation “boost” scores calculated above.

Not only do we find substantial variation between trinucleotides of the same class (S5 Fig), but we find the ranking within each GC class to be remarkably consistent between sequence types (5’ UTR, 3’ UTR, ncRNA, cis regulatory elements (CREs), introns) (Fig 8). We exclude coding sequence from this analysis to negate the impacts of coding selection. Within the most populated classes GC classes (33% and 66%), ranks are significantly correlated in all comparisons (Pearson’s method; all p < 0.01). This supports the hypothesis of a consistent isochore dependent fixation bias that acts differently on different trinucleotides of the same GC content. We note that the non-expressed CRE versus intron comparison gives exceptionally high repeatability indicating that transcription coupled repair/mutation probably does not explain these trends.

**Fig 8.**
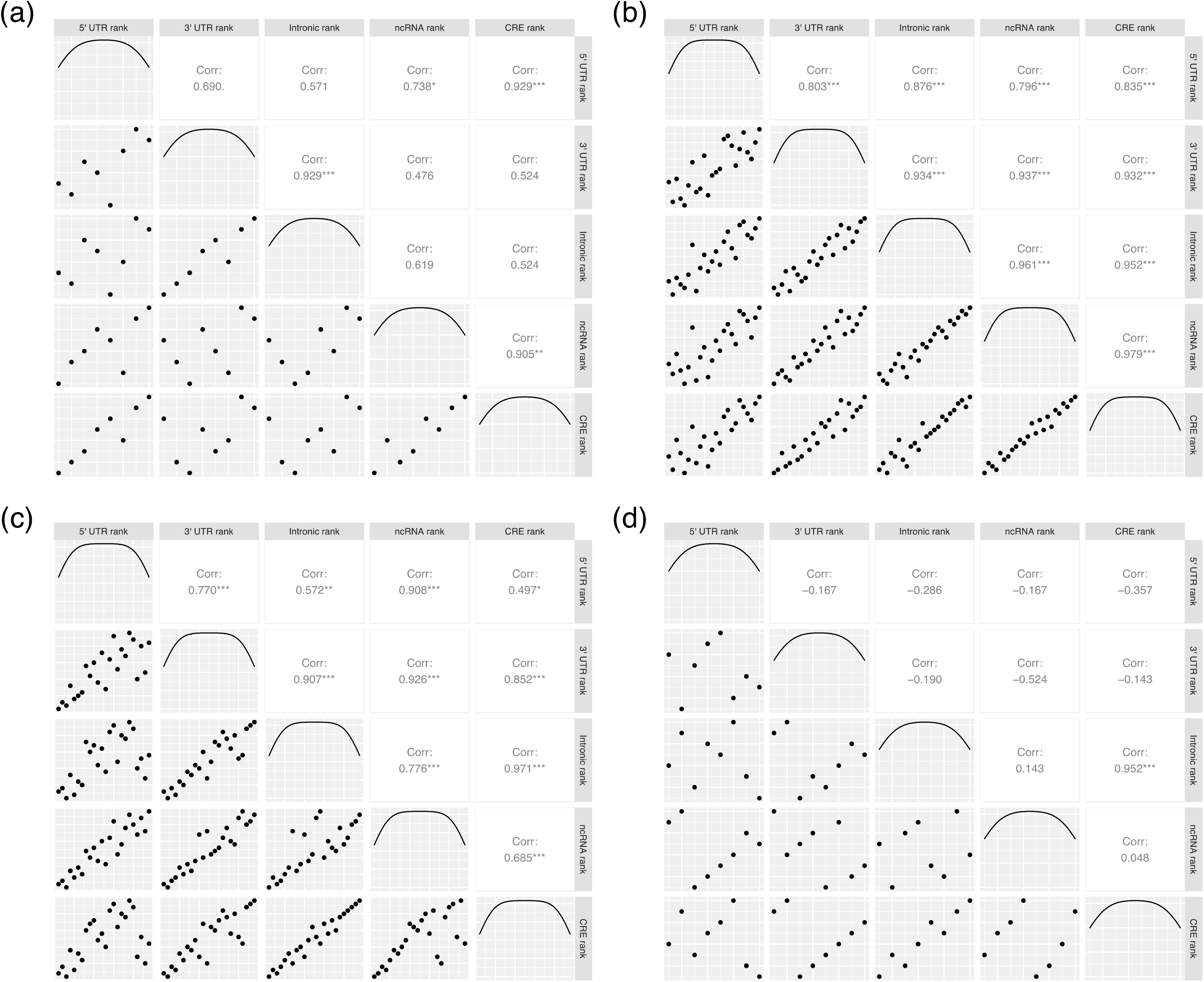
Correlation analysis of trinucleotide ranks (by their gBGC “boost” metric) within the four GC classes (a) 0%, (b) 33%, (c) 66%, (d) 100%. Within the 33% and 66% GC classes, ranks are significantly correlated in all comparisons (p < 0.01). This is not true of the 0% and 100% GC classes, correlation analyses within which are underpowered (n=8 trinucleotides in each class compared to 24 in the 33% and 66% classes). Correlation statistics were calculated using Pearson’s method.

Within the TAG/TGA case study, we find TAG to be less “boosted” than other A, G, T-containing trinucleotides, second only to GTA trinucleotides (S6a Fig). By contrast, TGA is the most promoted by fixation bias, with the one exception of AGT in the 5’ UTR (S6a Fig). Fixation bias correlated with GC-content hence appears to contribute to the differences in frequency between TGA and TAG trinucleotides outside of, and possibly also within, the stop codon context. That TCA also receives a consistent higher GC-coupled fixation boost than TAC (S6b Fig) favours that the fixation bias is dependent on nucleotide context rather than stop codon functionality. We also recall that while high recombination rate favours flux to the GC-rich state at two-fold degenerate sites, Glutamine (CAA→CAG) is one exception to this rule (S4 Fig). If CAA→CAG is suffering a similar fate to TAA→TAG this too would be supportive of a general nucleotide context-dependent trend in fixation bias affecting TAG rather than selection for termination efficiency.

## Discussion

The assumption that sequence conservation implies purifying selection and hence optimality of the preserved sequence underpins many enterprises, from medical diagnostics to evolutionary analyses of the proportion of sequence that is functional. While there has been prior consideration that tests for positive selection might be impacted by gBGC mimicking selection’s signatures (12, 35–40), there has been less attention paid to the problem that it might also explain sequence conservation, despite this being a logical necessity (41). We identified the case of stop codon usage in mammals as a test case because prior evidence suggested a contradiction: TAA looks to be optimal (as elsewhere) but TGA was nonetheless conserved. We reasoned that gBGC might explain this and resolve the exceptionalism of mammalian stop codon usage. Our data strongly support this. We see TGA usage is higher in GC-rich and highly recombinogenic domains, with the same trends also being seen in non-coding sequence. Increased TAA→TGA flux is also seen in GC rich regions and regions of high recombination. Multiple lines of evidence suggest that at the focal termination codon TGA is not optimal and hence that gBGC can act against the direction of selection. The results satisfy all criteria proposed by Duret and Galtier (11) for differentiating gBGC from selection. Across species a greater flux of TAA→TGA in the GC richer genes is associated with a greater intragenomic variance in GC content, consistent with the above trends being predicted, broadly speaking, by the extent to which a species is isochoric.

Is the TAA/TGA enigma a special case or indicative of a more general trend? We observe that deviation of all trinucleotides from mutational equilibrium in GC-rich domains is strongly predicted by their GC content. The TAA→TGA trend in high GC domains can be considered a special example. More generally then, we have strong reason to suspect the gBGC mediated fixation bias will cause false signals of purifying selection at GC-rich residues in GC-rich isochores that extend beyond the specific context of TAA→TGA flux. This example is however unusual in that we have confidence that the substitutional process at the focal termination codon context forces conservation of a non-optimal codon, a trend that can be partly overcome by stronger selection for optimality in highly expressed genes.

There is, however, another possibility to explain deviation from mutational equilibrium in domains of high GC, this being that some form of selection favours GC-rich sequence. As Hill-Robertson interference (104) is reduced in domains of high recombination selection should be more effective in such domains, causing a fixation bias. One can imagine many possible modes of such selection, for example on DNA structure (105–107) or on nucleosome positioning (108–111). Unlike gBGC that predicts GC enrichment, any selection model must, after the fact, explain why GC-rich trinucleotides are favoured. Such models are unconvincing for several reasons.

First, in the current context TGA is not selectively favourable at the focal termination codon but nonetheless conserved. This suggests we must evoke a force other than selection to explain TGA conservation (assuming selection on stop codon functionality to be the strongest mode of selection at the focal stop). Why we should not similarly evoke the same force outside of the termination context seems like special pleading. Second, that the GC biasing effect correlates with male not female recombination rates (22), suggests that the effects are not mediated by reduced Hill-Robertson interference (22).

Third, the strength of selection (and associated load) in species with low N_e_ (mammals and birds) is problematic. Consider the hypothesis that TA dinucleotides could lead to accidental incidences of, for example, “TATA” boxes in eukaryotes and “Pribnow” boxes in bacteria (i.e. the TATAA motif). More generally, TA features in many key regulatory motifs that would be inappropriate in most DNA regions in both eukaryotic and prokaryotic genomes (112, 113). To date this is probably the best (if not only) model for selection against TA in all taxa. This could, in principle, explain why TAG is underused compared with TGA. Indeed, within the trinucleotides with only A and T, ATA and TAT, the two that are core to TATA box, are consistently the two with the lowest “boost” (S6 Fig). In bacteria and archaea, the strength of selection against such spurious binding is estimated to be around *N*_e_*s* = -0.09 and thus within the range of nearly neutral mutations for these species (114). If then *Escherichia coli*’s N_e_ is of the order of 10^8^ (115), then *s* must be ∼-0.09/10^8^ = -9 x 10^-10^. For a mutation to be under selection in humans *s* ∼ 1/2 N_e_ must hold. In a species with N_e_ ∼ 10,000 (e.g. humans) then this value of *s* (i.e. 1/20,000) is much greater than 9 x 10^-10^ estimated for selection against spurious binding. Thus, unless the selective cost of spurious binding is very much greater in humans than in bacteria, it is hard to see how selection can be efficient enough to remove point mutations that introduce spurious binding sites.

We do not presume that mutation bias and selection have no role. Indeed, in GC-poor domains mutation bias appears to provide a robust fit to the observed trends and explains the differential usage of TAG and TGA. Further, highly expressed genes over-employ TAA. However, for a full explanation of TGA conservation, especially in GC-rich domains, we need to evoke some other force, of which biased gene conversion is a good possibility, not least because it predicts high GC trinucleotides should be given a fixation boost in GC-rich domains, as observed.

We do not wish to claim that TAA is optimal for all genes. There could be many reasons that, for some genes, TGA is optimal. One possibility could be that TGA might be the least leaky in some contexts but as the experimental evidence contradicts this possibility (61), we don’t consider this reasonable. Alternatively, TGA may be TR-prone and “leaky”, but that leakiness is selectively favoured in some instances. High rates of TR may beneficially increase proteome diversity (116). Indeed, a few examples of functional read-through have been described (117, 118), though the commonality of this in mammals is unknown. Alternatively, read-through may be part of a gene regulatory mechanism (76, 119). Indeed, the discovery of TGA conservation prompted speculation that TGA might be commonly optimal in humans as it enables novel gene expression control. Specifically, it was suggested that ribosomes that read through the primary stop codon stall and form a queue from the next in-frame stop (or ribosome pausing factor), filling the space between the two stops and eventually infringing upon the 3’ end of the coding sequence itself. At this point, translation of this mRNA molecule is blocked (119). The fact that readthrough occurs at a low (but not very low) rate thus allows the mRNA molecule to be translated a relatively tightly regulated number of times prior to degradation.

Generally, however, it is unclear how any adaptive TR model might explain mammalian exceptionalism in stop codon usage. Given that TGA optimality cannot explain why TGA is also favoured in non-canonical stop contexts, the above arguments are, by Occam’s razor, not needed to explain general trends. Moreover, were there selection for TR, one might expect this to be common to all eukaryotes and therefore predict higher TGA usage in species with high N_e_ (not just mammals), but this isn’t seen (94). Instead TAA usage correlates positively with N_e_ (94), as expected if it is the optimal stop codon (although there are mechanisms that are rare in high N_e_ species but common in mammals, a high density of exonic splice enhancers to define intron-exon junctions being a case in point (120)).

### Why do trinucleotides of the same nucleotide content have different fixation boosts?

Our evocation of gBGC to explain the general trends in GC rich domains is not a complete explanation. Importantly we see repeatable trends whereby GC-matched trinucleotides show consistent differences in levels of fixation bias “boost” in GC-rich isochores. For example, TAG is among the least “boosted” trinucleotides in the 33% GC class, compared to TGA which more highly exceeds its mutational equilibrium at high GC isochores. Similarly, TAG usage appears largely uncorrelated with local GC content. Any model (selection, mutation, or gene conversion) evoking a relationship between simple GC pressure and differences in nucleotide content cannot obviously account for a difference in boost between nucleotide matched trinucleotides (e.g. TAG and TGA).

Given the ability of our complex mutation bias model to predict trinucleotide usage in low GC domains (Fig 6), we assume that our mutation bias estimation in GC rich domains is also largely accurate. If so, complex mutation bias is unable to explain the repeatable boost scores (Fig 8). In principle there could be several remaining classes of explanation. First, selection might act differently on underlying di or trimers. For example, regarding TAG and TGA, selection on TA or AG residues may be different to that on TG or GA ones. We can find no convincing evidence for this that can explain the universality of TAG avoidance (see S1 Text). One also needs to evoke selection that is strong enough throughout the human genome, which appears unlikely for reasons given above.

A further possibility is an interaction between complex mutation bias and gBGC making certain trinucleotides more liable to conversion owing to their relative commonality in populations. With a difference in mutational equilibria, the incidence of TAA/TAG meiotic heteroduplex mismatches (or sense/antisense ones to be more precise) is highly likely to be lower than that of TAA/TGA mismatches. Thus, gBGC may more commonly act on TAA/TGA. Overall, however, we see no correlation between our gBGC boost score and mutational equilibrium in any GC class of trinucleotides (Spearman’s rank; p > 0.05 for 0%, 33%, 66%, and 100% GC trinucleotides). Pairwise comparison of all possible trinucleotide combinations also indicates that the trinucleotide with the higher mutational equilibrium does not necessarily receive the higher boost (Binomial test with null probability = 0.5, p = 0.17). This may reflect the fact that common trinucleotides are also more commonly substrates to be converted.

Finally, like mutation, gBGC may be contingent on the local sequence context such that, for example TAG and CAG are relatively unaffected by gBGC, while TGA is affected. This could explain similar trends in bacteria and eukaryotes if, as is claimed, gBGC also operates in bacteria (121). Complex specificity might be expected as many protein-nucleic acid interactions are contingent on local sequence context. For example, APOBEC3/A/B induced mutations account for many C→T and C→G mutations but occur predominantly in the context of TC[A/T] (122, 123). More specifically, several DNA repair processes are known to be affected by local sequence context (124) including, at least in bacteria, mismatch repair (125), the process underpinning gBGC. Here, sequence contexts that enhance localised DNA flexibility are associated with mismatch repair activation (125) (see also: 126, 127). Similar evidence for a role of local DNA flexibility has been found in yeast (127, 128). The biological response elicited by CTG and CGG repeats in human trinucleotide repeat disorders may be mediated by their increased flexibility indicative of a relationship between local flexibility and trinucleotide content (129). Evidence in humans for more effective repair of flexible DNA owing to local sequence context (130) suggests that an association between DNA mismatch repair and DNA flexibility may have relevance to understanding fixation biases in GC-rich domains. If flexibility is the core factor, then we might expect that a trinucleotide and its antisense should have similar boost scores as both feature in the same three base pairs of DNA (one on the Crick strand, the other on Watson). In our data, however, we find that the difference in gBGC “boost” between sense and antisense trinucleotides is no smaller than randomised trinucleotide comparisons (p > 0.05 regardless of the sequence analysed). This suggests that DNA flexibility alone cannot explain gBGC boost. Despite this, direct analysis of the sequence context associated with gBGC would be valuable.

## Methods

### General methods

All data manipulation was performed using bespoke Python 3.6 scripts. Statistical analyses and data visualisations were performed using R 3.3.3. All scripts required for replication of the described analyses can be found at https://github.com/ath32/gBGC. While stop codons function at the mRNA level, we here analyse chromosomal DNA sequences and therefore refer to the three stops as TAA, TGA and TAG.

### Inferring stop codon switches from eukaryotic triplets

Lists of one-to-one orthologous genes were downloaded for a diverse variety of species triplets from the main Ensembl repository (release 101), Ensembl plants (release 46), or Ensembl metazoan (release 46): (1) primates; *Homo sapiens, Otolemur garnettii, Callithrix jacchus,* (2) cows; *Bison bison bison, Bos grunniens, Bos taurus,* (3) dogs; *Canis lupus familiaris, Ursus americanus, Vulpes vulpes,* (4) mice/rodents; *Mus musculus, Mus spretus, Rattus norvegicus*, (5) birds; *Gallus gallus, Anas platyrhynchos platyrhynchos, Meleagris_gallopavo,* (6) flies; *Drosophila melanogaster, Drosophila pseudoobscura, Drosophila simulans*, (7) nematodes; *Caenorhabditis briggsae, Caenorhabditis remanei, Caenorhabditis elegans,* (8) plants; *Arabidopsis halleri, Arabidopsis lyrata, Arabidopsis thaliana*. Orthologous genes were extracted from the respective genomes using whole genome sequence and gene annotation data downloaded from the same sources. Genes were filtered to retain genes with CDS length divisible by 3, no premature stop codons, and stop codons TAA, TGA or TAG. Genes from each species triplet that met our quality controls were aligned using MAFFT with the -linsi algorithm (131).

Rather than using parsimony as done previously (49, 132), stop codon switches were reconstructed using a maximum likelihood approach. For each species triplet, ancestral nucleotide states for the internal node between the two ingroups were inferred by maximum likelihood using IQTree v2.1.2 with the -asr flag (133, 134). This analysis does have one limitation in that we do not control for the possibility of parallel substitutions, however we assume this effect to be small. To calculate stop codon flux rates, we compute the inferred ancestral stop codon state at the internal node and calculate transition from this ancestral state to a derived state (per incidence of the ancestral state).

### Predicting equilibrium TGA content using flux data

The predicted TGA usage for a given lineage, pTGA, was calculated by adapting the formulae outlined by Long, Sung (44). In their study, given a spectrum of de novo mutations, they propose the equilibrium GC content, P_n_, can be calculated from the GC→AT mutation rate divided by the reciprocal rate, m, such that:

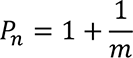

We adapt this equation to the stop codon exemplar. As TAA and TGA stop codon usage covary in opposite directions with genomic GC content we consider their usage to be dependent on one another. Due to the unusual biology of TAG, not least that it remains lowly used irrespective of genomic GC content, we exclude fluxes involving TAG from this calculation. Our proposed equation for calculating equilibrium TGA content, pTGA, from the ratio of TGA→TAA divided by TAA→TGA, s, is:

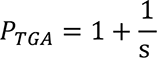

### Null simulations to assign significance to observed pTGA deviation between two groups of genes

The difference in pTGA observed between two gene groups (“A and B”, GC rich and GC poor genes, or highly recombining and lowly recombining genes) may be assigned significance by comparisons to simulated null gene groups. First, by analysing all genes en masse we can calculate a genomic rate of TAA→TGA per TAA and for TGA→TAA per TGA. For each group of genes, we may then calculate null pTGA scores that control for these rates.

For each gene in the group, we determine the ancestral stop codon (of which we are only interested in TAA or TGA) and record the number of each. If the ancestral stop codon is TAA we generate a random number between 0 and 1 and if equal to or below the genomic TAA→TGA rate we record a null TAA→TGA flux event. If the ancestral stop codon is TGA we generate a random number between 0 and 1 and if equal to or below the genomic TGA→TAA rate, we record a null TGA→TAA flux event. By this method we thus receive null counts of TAA→TGA and TGA→TAA which may be divided by the ancestral counts of TAA and TGA to receive null flux rates. From these rates we may calculate null pTGA, and thus by repeating this process 1,000 times we create a null distribution of pTGA for the gene group. Repeating this method for both gene groups, we have a distribution for gene group A and gene group B.

Next, we randomly sample with replacement one pTGA score from each of the two distributions, receiving a random pair. For each random pair we calculate the deviation between that sampled from group A and group B and repeat this process 10,000 times to create a null distribution of differences. We then compare the observed difference between the real gene groups to this distribution, asking how many simulants have as high a difference as the observed one (n). The significance of the observed difference beyond null may be represented as p = n / m where m is the number of random pairs considered.

### Intronic GC as a proxy for isochore GC content

Under the assumption that intronic GC reflects isochore GC content, intronic nucleotide sequences were extracted from one candidate genome within a species trio (e.g., the human genome was used as a representative of the primate triplet) using the appropriate GFF and WGS files downloaded from Ensembl (release 101). From the resulting spectrum of intronic GC contents, 10% percentiles were calculated, and genes were binned accordingly. The stop switch method described above was applied to each bin to measure changes in stop switch frequencies across intronic GC contents.

This binning method is effective to segregate genes evenly across intragenomic GC contents but does not allow comparisons between eukaryotic groups. To plot the stop codon switches of multiple different species on the same axis requires a GC-matching methodology. To achieve this, genes were binned at 5% intronic GC content intervals (e.g. genes of GC content between 27.5% and 32.5% would be allocated to the 30% bin). As this method does not use percentiles, the resulting bins are not pre-designated to be equal in size. Bins of insufficient size (n < 50) were discarded. As before, the stop switch method was then applied to the GC-matched bins to measure changes in stop switch frequencies.

### Calculating mutational equilibria

The equilibrium content of all four nucleotides (indicated N*) may be estimated using the full mutational spectrum (135, 136). A full spectrum of 108,778 de novo mutations (from 1,548 Icelandic human family trios) was downloaded from the supplementary material of Jonsson, Sulem (101). Knowing the rate of flux between every nucleotide (normalised to the occurrence of each nucleotide), we calculate the mutational equilibrium states of all nucleotides and GC content exactly as outlined in Rice at al. (2020). The same theory can be applied to the three stop codons to predict their equilibrium frequencies as follows, where TAA’ indicates the frequency of TAA after some period of time:

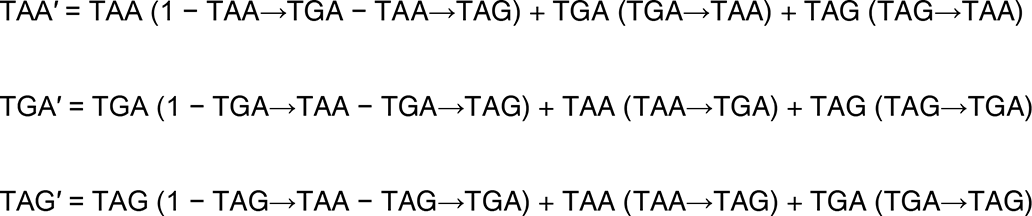

For equilibrium calculation, these simultaneous equations are solved such that TAA’ = TAA, etc. We are solving for gain = loss for each stop codon:

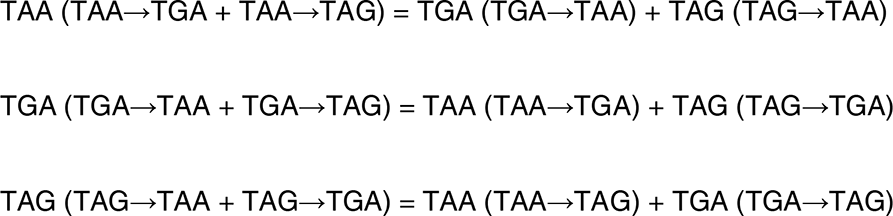

Note that in these equations we ignore the possibility of mutations from stop codons to sense codons. These we assume to be very rare and, should they occur, highly deleterious via the creation of C-terminal extensions. To constrain the results such that all equilibrium frequencies sum to 1, we replace one arbitrarily chosen stop codon frequency with 1 – the sum of the other two. While this would be achieved most accurately using precise mutational flux data between TAA, TGA, and TAG this is not captured within the Jonsson (101) dataset. Instead, we estimate flux between the three stops using null frequencies proposed by Belinky et al. (49). In their paper, they suggest the substitution control for TAA>TGA and TAA>TAG is A>G, for TGA>TAA and TAG>TAA is G>A, and for TGA>TAG and TAG>TGA is 2 x A>G x G>A.

The full spectrum of 108,778 de novo mutations may also be analysed using a 16×16 dinucleotide mutation matrix by tracing each mutation back to the reference genome and inferring dinucleotide changes. From the resultant matrix we estimate the equilibrium frequencies of each dinucleotide by adapting the simultaneous equations above to consider flux into and away from each dinucleotide. An estimated GC* may then be calculated from the 16 dinucleotide equilibria, whereas TGA* (and other trinucleotide equilibrium frequencies) may instead be estimated by incorporating the 16 equilibria into Markov models, simulating null sequences, and calculating trinucleotide frequencies from these (see “Markov models for simulating null sequences”).

### Gene expression metrics

To assess the role of gene expression in mammalian stop codon evolution we consider experimentally derived protein abundance data downloaded for *H. sapiens*, *B. taurus*, *C. familiaris*, and *M. musculus* from PaxDb (137). As selection acts on protein activity, not mRNA levels, we consider this a robust measure. For species where multiple datasets are available, we employ the whole organism integrated set for maximum coverage of the proteome (see https://github.com/ath32/gBGC for accessions list).

### Pseudo-autosomal regions, chromosome size, and local recombination rates

To assay the impact of recombination we employed a) chromosome size as a proxy of long-term recombination rate per bp, b) pseudoautosomal localization, this being known to be highly recombinogenic, and c) estimated recent recombination rates.

For the latter, we employed recombination rates generated by the HapMap2 project (138) using coordinates lifted to the hg19/GRCh37 human genome build by Adam Auton (available: ftp://ftp-trace.ncbi.nih.gov/1000genomes/ftp/technical/working/20110106_recombination_hotspots/). For this analysis we hence use the GRCh37 human genome build and annotations, downloaded from NCBI and available at: https://www.ncbi.nlm.nih.gov/assembly/GCF_000001405.13/ (last accessed 24 September 2020). For logistic regression modelling, each gene was assigned an estimated recombination rate equal to the average recombination rate of all its internal SNPs from the genetic map.

To assess the possible correlation between equilibrium GC content and recombination rate (see S7 Fig), we instead employ recombination rate bands directly assayed from 15,257 parent offspring pairs at 10kb resolution. This we consider to be the better data to use for this analysis as de novo mutations may be reasonably assigned the recombination rate of the 10kb band it falls within. The data were downloaded from https://www.decode.com/addendum/ (last accessed 14 September 2020) (139).

Coordinates of the two regions (PAR1 and PAR2) were downloaded from NCBI (https://www.ncbi.nlm.nih.gov/grc/human, last accessed 14 September 2020). Chromosome sizes employed are base pair lengths derived from human genome build hg38.

### Assessing the predictive abilities of gene expression and recombination rate

To determine whether expression and recombination rate can correctly predict the observed trends in stop codon usage we employ logistic regression. Stop codon usage and GC3 content was captured alongside gene expression data or recombination data (depending on the feature to be examined). Models were fit and examined using the glm function in R with the ‘family = binomial’ parameter. This produces a coefficient for each independent feature and associates a p-value for its predictive significance. We control for GC content by including GC3 content in a multivariate model when assessing expression level metrics. For the analysis of stop codon usage in null sequences we instead use linear regression, also using glm in R, as more than one ‘stop codon’ may be present in each sequence.

### PGLS analysis of TGA enrichment and effective population size (N_e_)

A phylogenetically-controlled test of correlation between N_e_ and TGA enrichment in lowly expressed genes (“LEGs”, lowest 25% of genes by protein abundance -see “gene expression metrics” above) were facilitated by PGLS using the “caper” R package (https://CRAN.R-project.org/package=caper). N_e_ estimates are from species with well resolved estimates of mutation rate and well described polymorphism data, and are the same as used in Ho and Hurst (94). Pagel’s lambda (λ) was predicted by maximum likelihood. Species used in this analysis were the same as published in our previous analysis (94), with the input phylogenetic trees generated using TimeTree (Kumar et al. 2017) and available in our GitHub repository along with the data required to repeat this analysis. TGA enrichment scores in LEGs were calculated such that:

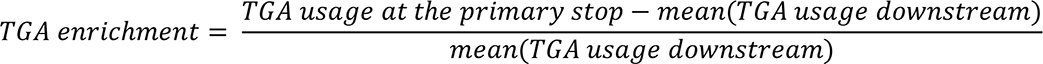

where mean TGA usage downstream is calculated from downstream codon positions +1 to +6. “Usage” refers to the relative frequency of TGA compared with the other stop codons TAA and TAA at position *n*, such that:

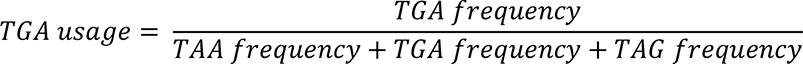

### Markov models for simulating null sequences

Null trinucleotide frequencies were generated from a null model that controls for underlying mono-or dinucleotide mutation rates. To achieve this, we first calculate mutational equilibrium frequencies for all mono- or dinucleotides - see “Calculating Mutational Equilibria” above and Rice, Morales (136). We next simulate 10,000 sequences (of average coding sequence length) using Markov models in a similar way to that outlined by Ho and Hurst (140). The first nucleotide/dinucleotide of each simulant is selected at random according to equilibrium nucleotide/dinucleotide frequencies. The following nucleotide is selected from a second set of frequencies: given the prior nucleotide in the simulation, what is the probability that the next nucleotide should be A, C, G or T. As all trinucleotides occur in these simulated sequences at a rate dictated by a derived mutational matrix, trinucleotide frequencies in the real sequences that are deviant from the simulations indicates enrichment or under-enrichment beyond chance.

## Data access

Raw data used are all publicly available and accessible as outlined in the methods section. All data manipulation was performed using bespoke Python 3.6 scripts. Statistical analyses and data visualisations were performed using R 3.3.3. Scripts required for replication of the described analyses can be found at https://github.com/ath32/gBGC.

## Acknowledgements

This work was supported by the European Research Council (grant EvoGenMed ERC-2014-ADG 669207 to L.D.H.).

